# The balance of local and distributed excitation shapes brain stability and reflects aging- and Alzheimer’s disease-related alterations

**DOI:** 10.64898/2026.09.17.752536

**Authors:** Yeongjun Park, Sunghun Kim, Hyunjin Park, Theodore D. Satterthwaite, Boris C. Bernhardt, Bo-yong Park

## Abstract

Human brain function emerges from the interplay between recurrent local activity and distributed inter-regional interactions. However, a biologically interpretable framework for quantifying this balance between the two factors at the whole-brain level remains lacking. Here, we extended an established biophysical model to quantify inter-regional excitation and newly introduced the recurrent ratio (R-ratio), a measure of the relative balance between intra- and inter-regional excitation. Dynamical analyses showed that an optimal R-ratio supports a trade-off between network stability and flexibility. Applying this framework to healthy aging and Alzheimer’s disease revealed progressively increased R-ratios with advancing age and disease severity. In healthy individuals, higher R-ratios were associated with age-related alterations in brain morphology, molecular pathology, and cognitive function, whereas Alzheimer’s disease was characterized by increased R-ratios accompanied by reduced dynamical persistence. Together, these findings establish the R-ratio as a biologically interpretable marker of whole-brain excitation balance and demonstrate its utility for linking biophysical mechanisms with large-scale brain dynamics, aging, and neurodegeneration.

## INTRODUCTION

Balance is a fundamental organizing principle of complex systems. Predator-prey interactions regulate ecological dynamics, whereas supply-demand relationships stabilize economic systems. Similarly, human brain function emerges through balanced interactions across multiple spatial scales. At the local circuit level, neural function depends on mechanisms such as competition between neural states during decision-making and excitation-inhibition balance for maintaining stable neuronal activity (Rubin et al., 2017; Wang, 2002). At the systems level, brain function emerges from the relative balance between intra-regional specialization and inter-regional integration across distributed neural networks (Arvin et al., 2022; Betzel and Bassett, 2018; Deco et al., 2021). Therefore, understanding how these balances are established and maintained is fundamental to explaining large-scale brain function.

The relative contributions of intra- and inter-regional processing influence multiple aspects of brain function. From an information processing perspective, they determine the trade-off between maintaining existing information and integrating new information (Shine and Poldrack, 2018; Wibral et al., 2014). Previous studies have suggested that transmodal regions support information maintenance through recurrent excitatory dynamics, whereas unimodal regions are more closely related to processing external inputs (Murray et al., 2014; Wang, 2001). They also govern the stability and flexibility of large-scale brain dynamics, enabling transitions between neural states that support adaptive cognition (Hancock et al., 2024; Shine et al., 2019). Disruptions of this balance are associated with aging and neurological disorders. Aging is associated with reduced long-range connectivity, potentially increasing reliance on local processing (Geerligs et al., 2015). Similarly, Alzheimer’s disease (AD) is characterized by accumulation of amyloid-*β* and tau, which may disrupt large-scale brain connectivity (Guzmán-Vélez et al., 2022; Wales and Leung, 2021). Together, these connectivity alterations may shift the balance between intra- and inter-regional interactions. Despite its central role in brain organization, quantitative frameworks directly characterizing the relative contributions of intra- and inter-regional processing within a biologically plausible model have only been sparsely explored.

Several computational approaches have been proposed to characterize neural dynamics. Effective connectivity models estimate directed interactions among brain regions, but their application to whole-brain dynamics remains limited by the temporal resolution of neuroimaging data and computational complexity (Friston et al., 2014, 2003). State-space models can dissociate recurrent and input-driven components while capturing temporal dynamics, but the biological interpretation of their latent states often remains ambiguous (Pandarinath et al., 2018; Vidaurre et al., 2017). Information-theoretic approaches quantify local information storage and distributed information transfer but provide limited mechanistic insights into the underlying neural circuitry (Vicente et al., 2011; Wibral et al., 2014). Collectively, these limitations highlight the need for a biologically interpretable framework that supports whole-brain simulations, captures temporal dynamics, incorporates recurrent processing, and provides mechanistic insights into large-scale brain function.

Large-scale circuit models provide such a framework by extending biologically plausible local circuit models to whole-brain networks through mean-field approximations (Deco et al., 2013a; Wang et al., 2019; Wong and Wang, 2006). Previous studies have used these models to investigate metastability, synchronization, and attractor dynamics (Demirtaş et al., 2019; Hancock et al., 2024). They have also been applied to characterize disease-related alterations in brain dynamics (Demirtaş et al., 2017; Park et al., 2021; Patow et al., 2023; Weng et al., 2020) and to investigate cognitive functions and their neurobiological substrates (Kong et al., 2021; Saberi et al., 2025). However, these studies have primarily focused on reproducing large-scale brain activity and its emergent dynamics, rather than quantifying how the balance between intra- and inter-regional contributions shapes whole-brain dynamics and function.

To fill the gap, we extended the parametric feedback inhibition control (pFIC) model (Deco et al., 2014; Zhang et al., 2024) to derive a quantitative measure of the relative contributions of intra- and inter-regional interactions, termed the recurrent-ratio (R-ratio). By computing the R-ratio, we established the relationship between the R-ratio and the excitation/inhibition (E/I) ratio to demonstrate whether the R-ratio captures a biologically meaningful property of local circuit organization. Next, we investigated how variations in the R-ratio shape large-scale brain dynamics. Finally, we evaluated how the R-ratio evolves in both healthy aging and neurodegeneration. As described below, our results provide a biologically interpretable framework for linking local circuit properties to large-scale brain dynamics and function.

## RESULTS

### Study overview

We leveraged multimodal magnetic resonance imaging (MRI) data from 1,004 neurologically healthy participants in the Human Connectome Project (HCP; Van Essen et al., 2013). To investigate the characteristics of intra- and inter-regional contributions to large-scale brain dynamics, we first derived the R-ratio, a quantitative measure of the relative balance between intra-regional recurrent excitation and inter-regional excitatory amplification, by optimizing modified biophysical models (**Fig. 1A**). We then characterized the mechanistic properties of the R-ratio by examining its relationships with the E/I ratio and large-scale dynamical responsiveness (**Fig. 1B**). Finally, we evaluated the clinical relevance of the R-ratio by investigating its alterations during normal aging and across the AD continuum using data from the Alzheimer’s Disease Neuroimaging Initiative (ADNI; Weiner et al., 2012; **Fig. 1C**).

**Fig. 1.**
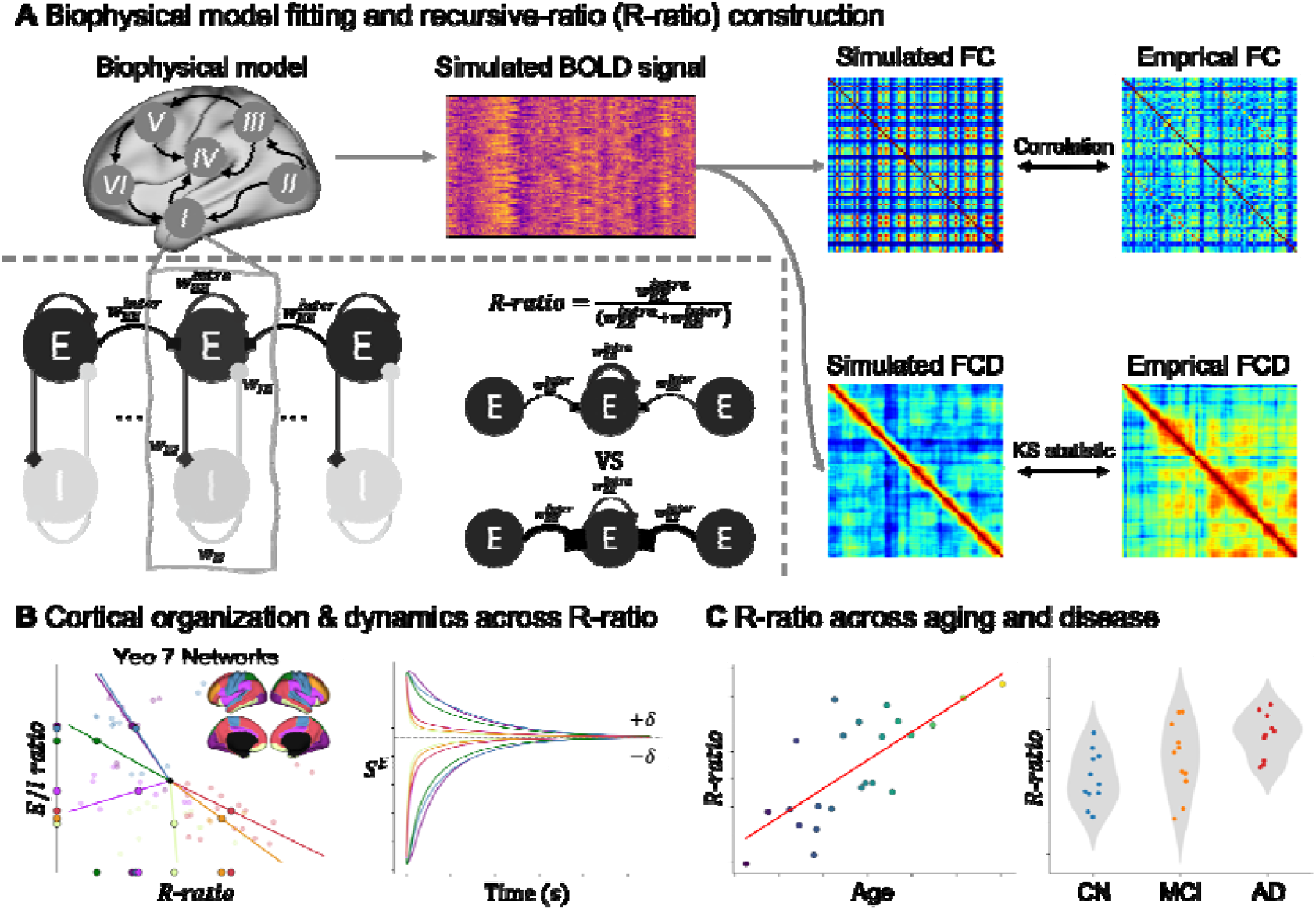
Schematic workflow for computing the R-ratio and its mechanistic interpretation. **(A)** The biophysical model was optimized to reproduce empirical FC and FCD, and the R-ratio was derived from the fitted control parameters to quantify the balance between intra- and inter-regional contributions. **(B)** The relationship between the R-ratio and the E/I ratio is illustrated across seven intrinsic functional networks (left). Dynamic analyses were performed to characterize the relationship between the changes in the R-ratio and model stability. **(C)** Age- and disease-related alterations in the R-ratio are explored. *Abbreviations: AD, Alzheimer’s disease; BOLD, blood-oxygen-level-dependent; CN, cognitively normal; E/I, excitation/inhibition; FC, functional connectivity; FCD, functional connectivity dynamics; KS, Kolmogorov–Smirnov; MCI, mild cognitive impairment*.

### Biophysical model optimization

We modified the established pFIC model to quantify intra- and inter-regional excitatory contributions to regional neural dynamics. In the original pFIC model, local recurrent excitation is controlled by the excitatory connection strength, whereas inter-regional excitation is transmitted through the structural connectivity (SC) and uniformly scaled across the cortex by a global coupling parameter. This formulation assumes that the scaling of long-range excitatory input is spatially homogeneous and therefore cannot quantify region-specific differences in inter-regional excitatory contributions. To overcome this limitation, we relaxed the global scalar parameter with a region-specific term to capture inter-regional excitatory interactions. The original recurrent excitatory parameter was reformulated as the intra-regional excitatory connection strength and inter-regional excitatory connection strength (see *Methods* for details).

We optimized this new model using loss functions based on functional connectivity (FC) and FC dynamics (FCD) (Kong et al., 2021; Zhang et al., 2024). The modified model reproduced empirical whole-brain dynamics with performance comparable to that of the original pFIC model. The Pearson correlation between simulated and empirical FC was r = 0.71 for the modified model and r = 0.72 for the original pFIC model (**Fig. 2A**). The modified model showed a lower L1 distance between simulated and empirical FC matrices than the original model (ours: d = 0.09; pFIC: d = 0.11; **Fig. 2B**). Both models yielded the same Kolmogorov–Smirnov (KS) distance between empirical and simulated FCD distributions (KS = 0.15; **Fig. 2C**). Finally, the modified model yielded a slightly lower total loss than the original pFIC model across the top 10 parameter sets from the test dataset (ours: 0.544 ± 0.009; pFIC: 0.554 ± 0.011; p = 0.054; **Fig. 2D**). These results indicate that introducing region-specific inter-regional excitatory coupling preserved the ability of the pFIC model to reproduce empirical static and dynamic FC.

**Fig. 2.**
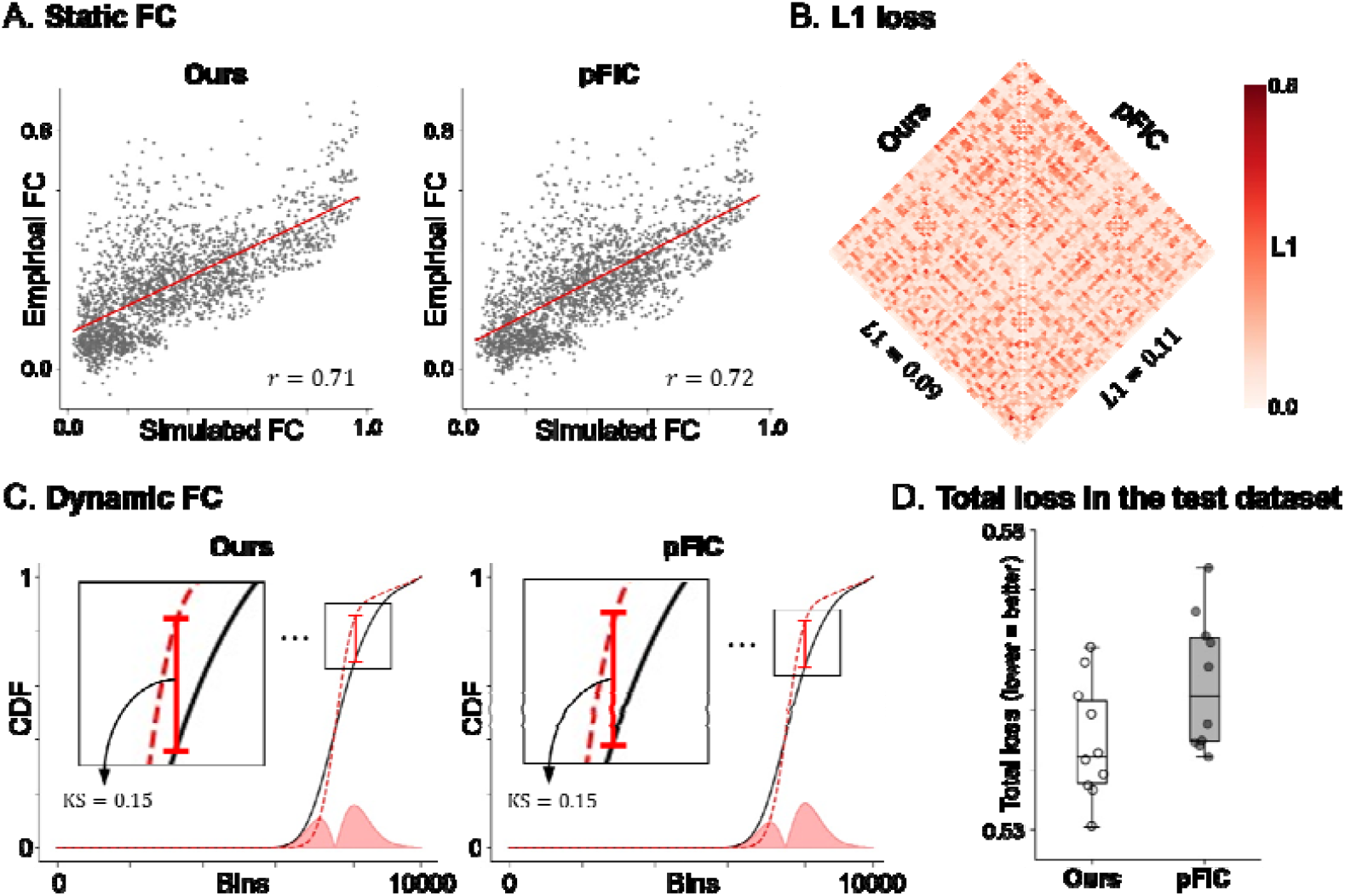
Performance of the modified pFIC model. **(A)** Pearson correlations between simulated and empirical FC matrices for the modified model (left) and the original pFIC model (right). **(B)** L1 distances between simulated and empirical FC matrices for the two models. **(C)** KS statistics between empirical (black line) and simulated FCD (red line) distributions. **(D)** Total loss evaluated on the test dataset. *Abbreviations: CDF, cumulative distribution function; FC, functional connectivity; FCD, functional connectivity dynamics; KS, Kolmogorov–Smirnov; pFIC, parametric feedback inhibition control*.

### Regional balance between intra- and inter-regional excitation

The modified pFIC model revealed distinct spatial distributions of the fitted control parameters (**Fig. 3A-C**). The inter-regional excitatory parameter () was highest in unimodal regions, including the visual and somatomotor cortices, and progressively decreased toward transmodal association cortices (**Fig. 3A**). In contrast, excitatory-to-inhibitory connection strength () exhibited the opposite pattern, with higher values in transmodal regions and lower values in unimodal regions (**Fig. 3C**). The intra-regional excitatory parameter 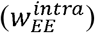 showed relatively modest regional variation (**Fig. 3B**). To determine whether the 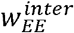 reflects underlying cortical microstructure, we examined the relationship between the optimal 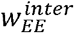 in the test dataset and microstructural profile (MP) moments derived from T1w/T2w imaging (**Fig. 3D**). The optimal 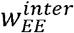 showed significant positive correlations with MP mean (r = 0.69 and p_spin_ < 0.001) and standard deviation (r = 0.52 and p_spin_ < 0.001), whereas significant negative correlations were observed with MP skewness (r = −0.71 and p_spin_ < 0.001) and kurtosis (r = −0.68 and p_spin_ < 0.001). These findings indicate that regional differences in inter-regional excitatory gain are systematically associated with cortical microstructural organization.

**Fig. 3.**
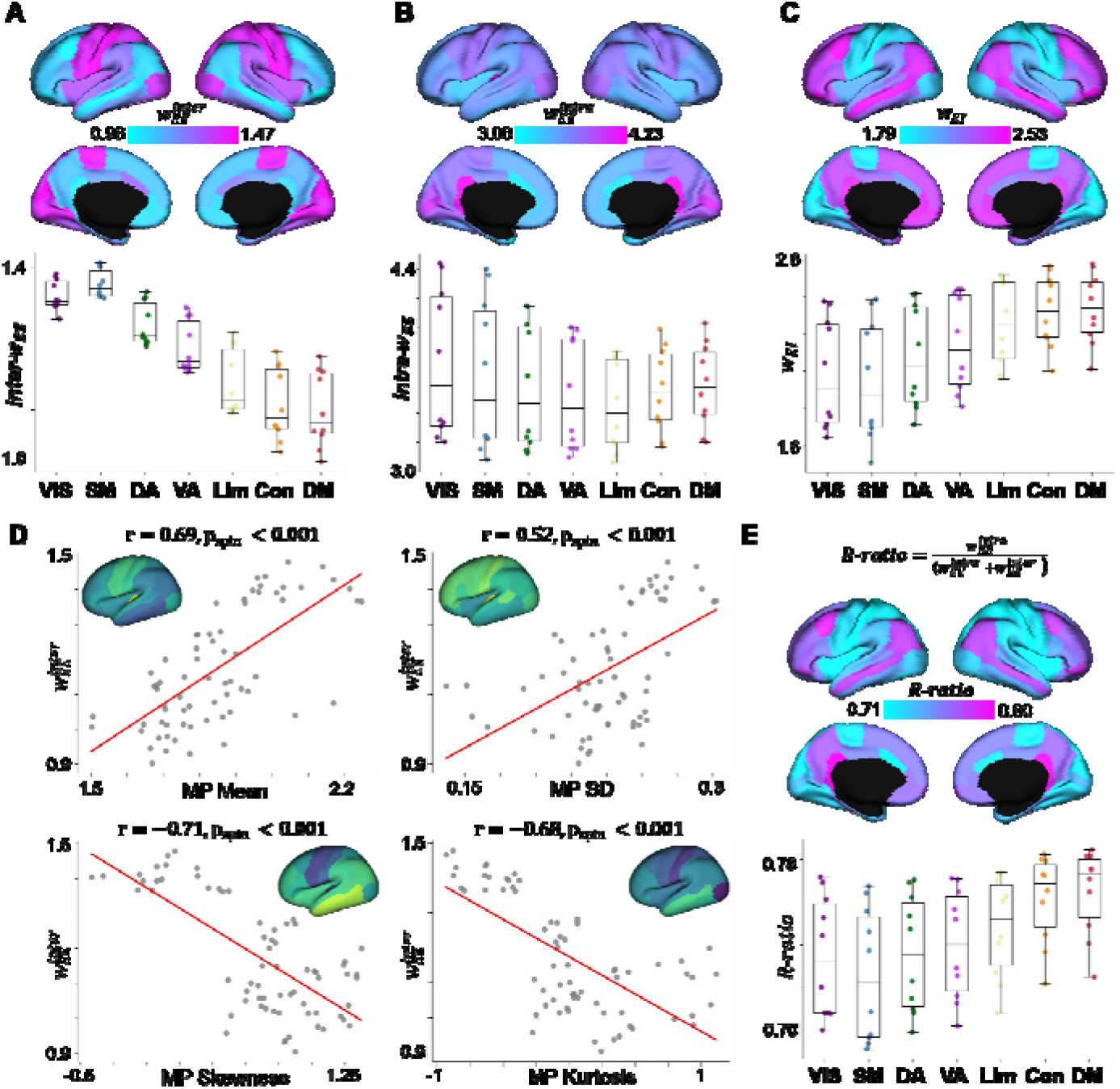
Spatial organization of intra- and inter-regional excitatory parameters and the R-ratio. **(A)** Spatial distributions of the, **(B)**, and **(C)** are shown on cortical surfaces (top). Parameter values are stratified across intrinsic functional networks (bottom). Box plots show the interquartile range, with center lines indicating median values. **(D)** Spatial correlations between and MP moments, including mean, standard deviation (SD), skewness, and kurtosis. **(E)** Spatial distribution of the R-ratio across the cortex (top) and intrinsic functional networks (bottom). *Abbreviations: VIS, visual; SM, somatomotor; DA, dorsal attention; VA, ventral attention; Lim, limbic; Con, frontoparietal control; DM, default mode; MP, microstructural profile; R-ratio, recurrent-ratio*.

Based on the distinct functional properties of unimodal and transmodal regions (Murray et al., 2014; Wang, 2001), we hypothesized that the relative balance between intra-regional recurrent excitation and inter-regional excitatory amplification would vary along cortical hierarchy. To quantify this balance, we defined the R-ratio for the *i*-th region as 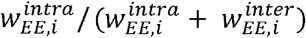. This measure reflects whether regional excitatory activity is more strongly shaped by intra-regional recurrent excitation or inter-regional excitatory amplification. The R-ratio was lowest in unimodal regions and progressively increased toward transmodal association regions (**Fig. 3E**). Across intrinsic functional networks, sensory networks exhibited the lowest R-ratio values, whereas frontoparietal control and default mode networks showed the highest values. The primary findings were based on the Desikan–Killiany atlas, and these spatial patterns were replicated using the Schaefer 100 atlas (**Supplementary Fig. 1A**). These findings suggest that the balance between recurrent and distributed excitation follows the canonical sensory-to-association cortical hierarchy, with unimodal regions relying predominantly on distributed excitatory inputs and transmodal regions exhibiting stronger recurrent excitatory dominance.

### The space defined by the R-ratio and E/I ratio captures multimodal cortical architecture

To characterize cortical regions in terms of both the source of excitatory drive and E/I ratio, we constructed a two-dimensional feature space defined by the optimal R-ratio and E/I ratio in the test dataset. Each cortical region was projected onto this space according to its R-ratio and E/I ratio, and the mean value of each measure was used to divide the space into four quadrants (**Fig. 4A**). Notably, only a small number of regions occupied the quadrant characterized by both high R-ratio and high E/I ratio, indicating that strong recurrent excitation rarely coexists with high excitatory dominance across the cortex. We next examined how intrinsic functional networks were organized within this two-dimensional space (Yeo et al., 2011) (**Fig. 4B**). The combined R-ratio and E/I ratio revealed a structured organization of intrinsic functional networks, forming a counterclockwise trajectory from sensory to association systems across the two-dimensional space (**Supplementary Fig. 2**).

**Fig. 4.**
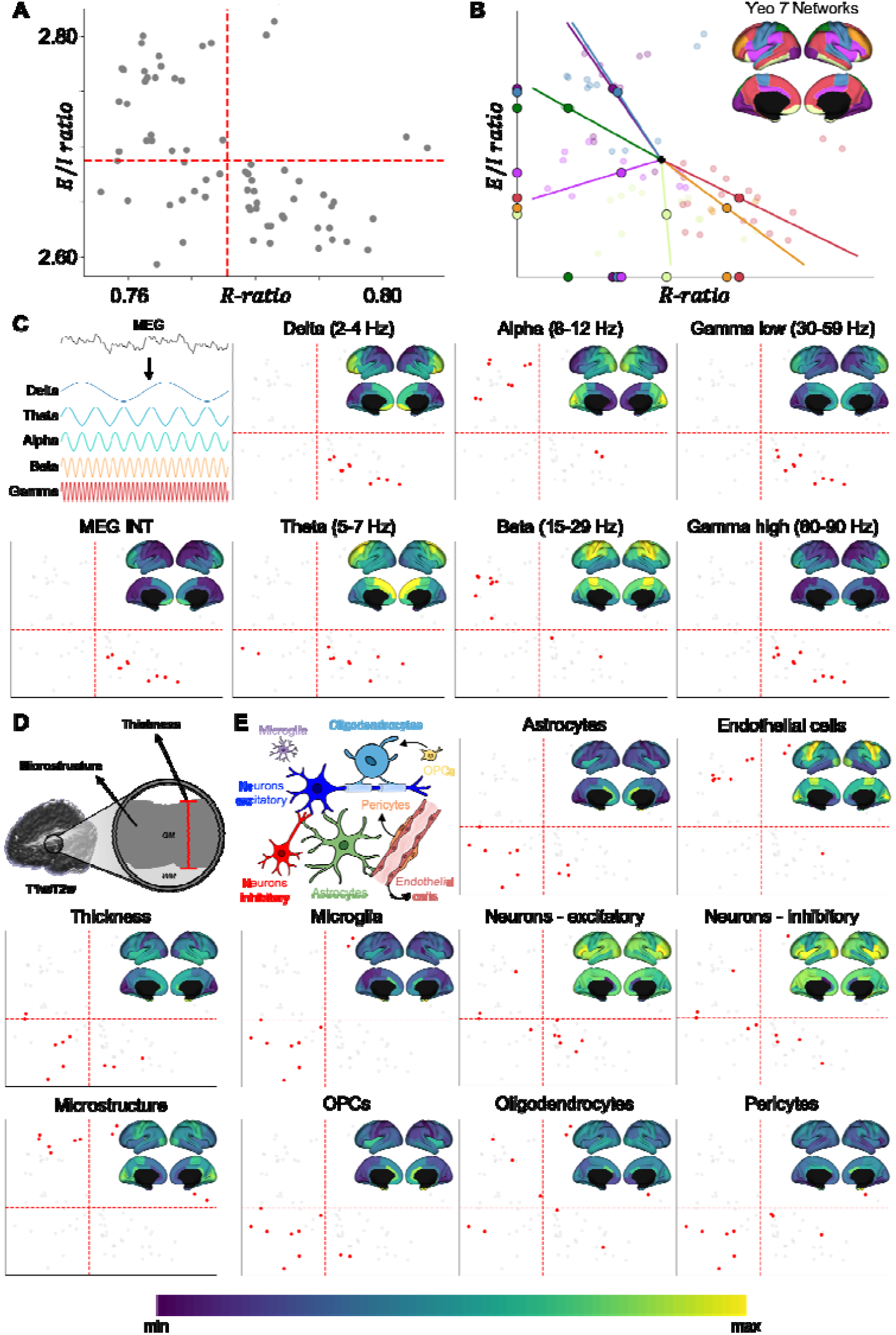
Multimodal contextualization of the R-ratio and E/I ratio. **(A)** Distribution of cortical regions within the two-dimensional space defined by the R-ratio and E/I ratio. Red lines indicate the mean values of each metric. **(B)** Distribution of the seven intrinsic functional networks within the R-ratio and E/I ratio space. The black cross indicates the mean R-ratio and E/I ratio values, and enlarged markers indicate the centroid of each network. The colored lines connect the black cross to the centroid of each network. **(C)** Distributions of cortical regions exhibiting the highest 10% of MEG-derived INT and frequency-specific power across the R-ratio and E/I ratio space. **(D)** Distribution of cortical thickness and T1w/T2w-derived myelin-related contrast. **(E)** Distribution of cell-type-specific gene expression profiles. Regions with the top 10% of each feature are highlighted in red. *Abbreviations: E/I, excitation/inhibition; MEG, magnetoencephalography; INT, intrinsic neural timescale; T1w, T1-weighted; T2w, T2-weighted; OPCs, oligodendrocyte precursor cells*.

To further characterize this organizational framework, we incorporated magnetoencephalography (MEG)-derived intrinsic neural timescale (INT) and frequency-specific power distributions (Markello et al., 2022; Van Essen et al., 2013). The top 10 regions with the highest values of each MEG-derived feature were highlighted in the two-dimensional space (**Fig. 4C**). Regions with long INT were predominantly located in the quadrant characterized by high R-ratio and low E/I ratio. Across frequency bands, regions with high alpha- and beta-band power primarily occupied the quadrant with low R-ratio and high E/I ratio, whereas regions with high delta-, theta-, and gamma-band power were concentrated in the quadrant with high R-ratio and low E/I ratio. We further assessed the organization of the R-ratio and E/I ratio space in relation to structural and molecular properties of the cortex (**Fig. 4D-E**). Regions with greater cortical thickness were predominantly located in quadrants with lower E/I ratios, whereas highly myelinated regions preferentially occupied quadrants with higher E/I ratios (**Fig. 4D**). Cell-type-specific gene expression profiles further exhibited that astrocytes, microglia, oligodendrocyte precursor cells, and pericytes were enriched in regions with both low R-ratio and low E/I ratio, whereas endothelial cell expression was concentrated in regions with low R-ratio and high E/I ratio (**Fig. 4E**). In contrast, excitatory neurons, inhibitory neurons, and oligodendrocytes were aligned along a linear axis within the two-dimensional space, suggesting coordinated variation of the R-ratio and E/I ratio across these cell populations. Together, these findings demonstrate that the R-ratio, when interpreted jointly with the E/I ratio, captures multiple organizational principles of the human cortex, spanning functional hierarchy, intrinsic electrophysiological dynamics, cortical microstructure, and molecular architecture.

### R-ratio shapes whole-brain dynamical properties

We next investigated whether variations in the R-ratio give rise to distinct whole-brain dynamical properties. To this end, we independently varied the internal scaling factors *β*_1_ and *β* applied to the optimal 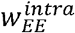 and 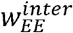 in the test dataset, respectively (see *Methods* for details), thereby modulating the balance between intra- and inter-regional excitation (**Fig. 5A**). We first performed a bifurcation analysis across the parameter space to identify transitions between stable and unstable dynamical regimes (**Supplementary Fig. 3**). We then evaluated how well each parameter combination reproduced empirical brain dynamics by computing the model loss on the test dataset (**Fig. 5B** and **Supplementary Fig. 4**). The optimal parameter combination was located close to the bifurcation boundary, which separates stable and unstable regimes of model dynamics. Parameter regions with either excessively high or low R-ratio values exhibited substantially larger loss values than the optimum, indicating that both extremes deviated from empirical brain dynamics. These findings suggest that whole-brain dynamics are best reproduced within an intermediate regime that balances intra- and inter-regional excitatory interactions.

**Fig. 5.**
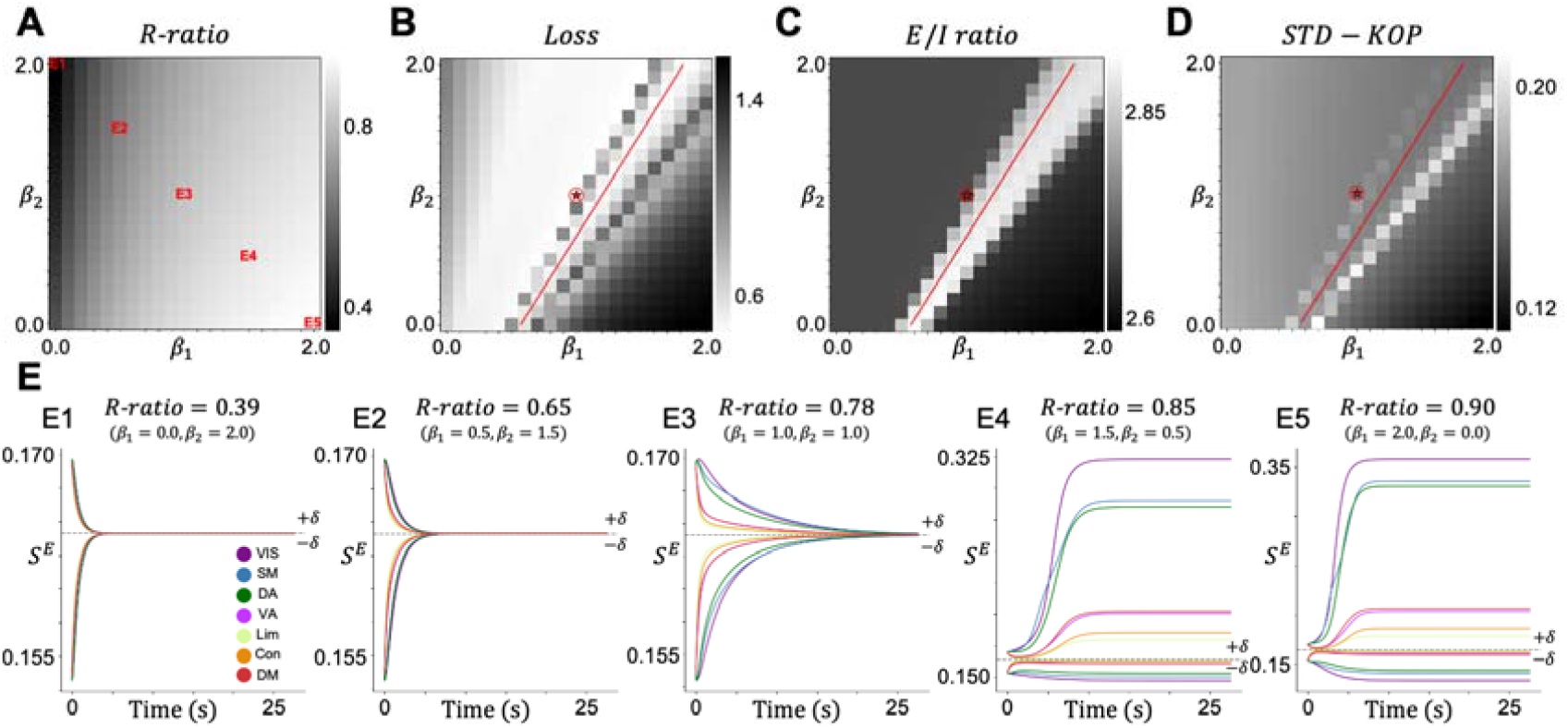
Whole-brain dynamical properties across the R-ratio parameter space. **(A)** Parameter space generated by independently varying the scaling factors applied to the intra-regional 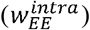 and inter-regional 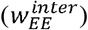 excitatory parameters. **(B-D)** Distributions of the model test loss, E/I ratio, and STD-KOP across the parameter space. The optimal parameter combination is indicated by the red circle with a star, and the red line denotes the bifurcation boundary separating stable and unstable dynamical regimes. **(E)** Representative perturbation responses across five R-ratio values (E1–E5). Each curve represents the mean synaptic activity (S^E^) averaged within each intrinsic functional network, with different colors indicating different networks. *Abbreviations: E/I, excitation/inhibition; STD-KOP, standard deviation of the Kuramoto order parameter*.

Next, we examined how variations in the R-ratio influenced intrinsic dynamical properties using the E/I ratio and the standard deviation of the Kuramoto order parameter (STD-KOP). The E/I ratio was used to assess the balance between excitatory and inhibitory dynamics, whereas STD-KOP was used to quantify temporal variability in global synchronization, which can be interpreted as a proxy for metastability (Hancock et al., 2024; Kuramoto, 1984). At the optimal parameter combination, the E/I ratio remained at an intermediate level, indicating balanced excitatory dynamics (**Fig. 5C**). In contrast, parameter regions near the bifurcation boundary exhibited markedly elevated E/I ratios, reflecting increased excitatory dominance. Similarly, the STD-KOP also exhibited intermediate values at the optimal parameter combination, indicating balanced synchronization dynamics (**Fig. 5D**). Regions near the bifurcation boundary showed substantially higher STD-KOP values, suggesting greater temporal fluctuations in large-scale synchronization.

Finally, we investigated how R-ratio shapes dynamical responsiveness by applying external perturbations (*δ*) around the fixed point and tracking the subsequent evolution of synaptic gating activity (**Fig. 5E**; see *Methods* for details). Parameter regimes with lower R-ratio values exhibited rapidly converging trajectories following perturbation, with short activity persistence times (**E1**: 1.22 s; **E2**: 1.86 s; **E3**: 7.25 s; **Fig. 5E1-3**). In contrast, higher R-ratio regimes showed diverging trajectories (**Fig. 5E4-5**). Thus, lower R-ratio values were associated with rapid attenuation of perturbation-induced activity, whereas higher R-ratio values exhibited increased sensitivity to perturbations and reduced dynamical stability.

### R-ratio captures age-related differences in brain structure and function

To evaluate age-related effects of the R-ratio, we examined its associations with amyloid burden, brain morphology, and neurocognitive performance using data from multiple independent datasets (Jaffe et al., 2022; Park et al., 2025; Weiner et al., 2012) (**Fig. 6A**). We first estimated the R-ratio across age using cognitively normal (CN) participants (imaging sessions = 689) from the ADNI dataset (Weiner et al., 2012). Participants were sorted by age and divided into age bins of 30 participants each. Within each age bin, 15 participants were randomly selected for the training set, while the remaining participants were assigned to the test set. The optimal global R-ratio in the test set progressively increased with age (r = 0.74, p < 0.001; **Fig. 6B** and **Supplementary Fig. 1B**). At the molecular level, age-related increases in the global R-ratio were positively associated with amyloid burden (r = 0.47, p = 0.023; **Fig. 6C**). We next examined associations with structural brain measures. The global R-ratio was negatively correlated with cortical thickness (r = −0.67, p < 0.001), gray matter volume (r = −0.75, p < 0.001), and white matter volume (r = −0.74, p < 0.001) (**Fig. 6D-E**). In contrast, ventricular volume showed a significant positive correlation with global R-ratio (r = 0.73, p < 0.001; **Fig. 6E**). These findings suggest that higher R-ratio values are associated with age-related structural brain alterations. To assess the functional relevance of the R-ratio, we further examined its associations with neurocognitive performance across four cognitive domains, including executive function, memory, visual attention, and reasoning (see *Methods* for details). Significant positive correlations were observed across all domains, indicating that higher R-ratio values were associated with poorer cognitive performance. Together, these findings demonstrate that the R-ratio captures coordinated age-related alterations spanning molecular pathology, brain structure, and cognitive function.

**Fig. 6.**
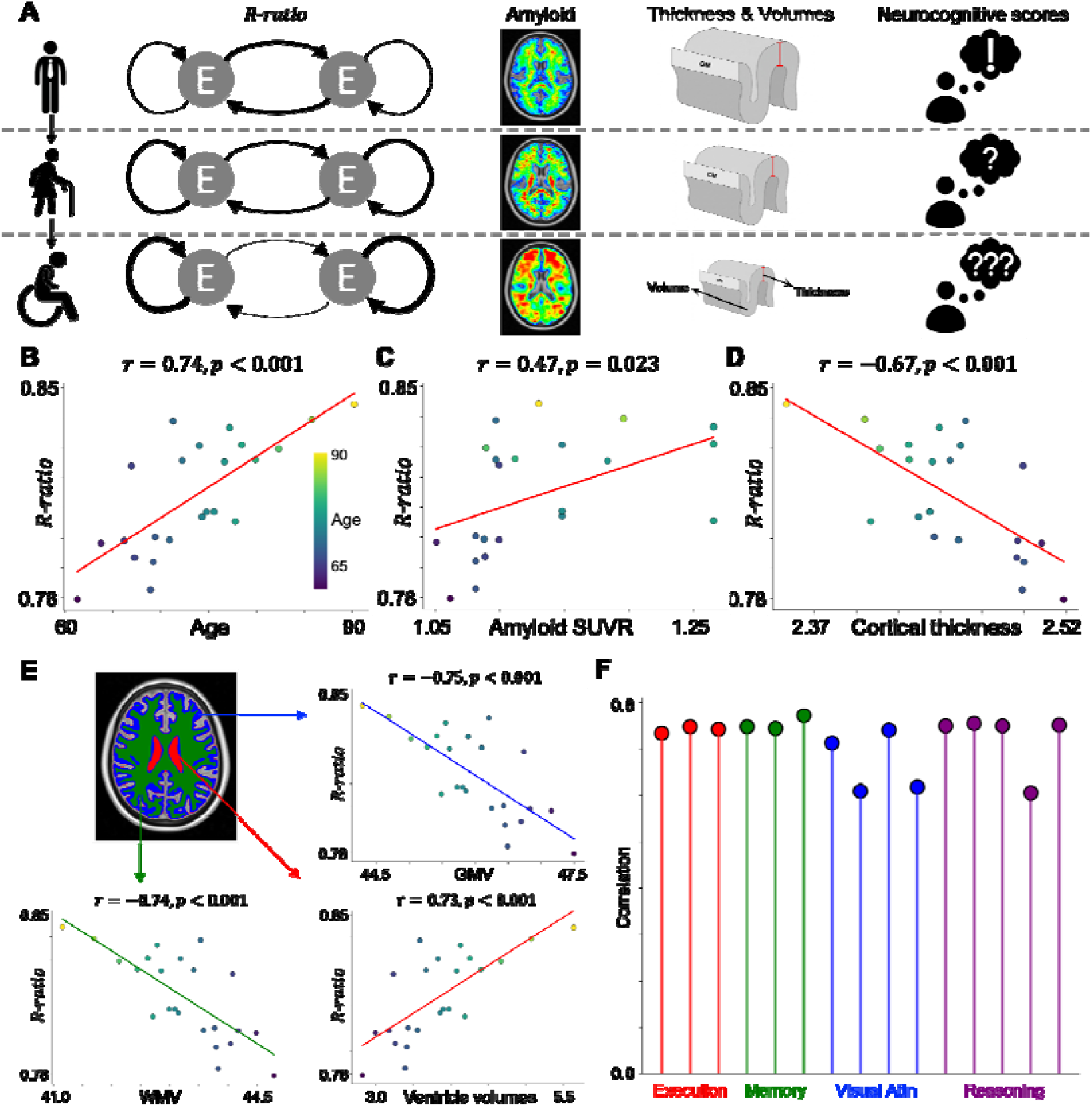
Structural and functional correlates of age-related differences in the R-ratio. **(A)** Schematic illustration of age-related differences in the R-ratio, amyloid burden, brain morphology, and neurocognitive performance. **(B)** Age-related differences in the R-ratio. **(C)** Correlations between age-related differences in the R-ratio and amyloid SUVR, **(D)** cortical thickness, and **(E)** gray matter, white matter, and ventricular volumes. **(F)** Correlations between age-related differences in the R-ratio and neurocognitive performance. *Abbreviations: SUVR, standardized uptake value ratio; GMV, gray matter volume; WMV, white matter volume; Visual Attn, visual attention*.

### R-ratio alterations across the Alzheimer’s disease continuum

Because the R-ratio was associated with age-related alterations in brain structure and function, we next investigated whether it also captures disease progression across the AD continuum. We analyzed phenotypic and imaging data from CN (imaging sessions = 689), mild cognitive impairment (MCI; imaging sessions = 436), and AD (imaging sessions = 137) participants from the ADNI dataset (Weiner et al., 2012).

Age-matched diagnostic groups were constructed in 2-year age intervals between 60 and 80 years, with 30 participants selected from each diagnostic group within each interval (**Fig. 7A**). Within each group, 15 participants were randomly selected as the training set, while the remaining participants were assigned to the test set. The optimal global R-ratio in the test dataset showed a progressive increase across the AD continuum, with significantly higher values in the AD group than in the CN group (p < 0.001; **Fig. 7B** and **Supplementary Fig. 1C**). We next projected the diagnostic groups onto the two-dimensional R-ratio-to-E/I ratio space. Disease progression was accompanied by a shift toward regions characterized by higher R-ratio and lower E/I ratio values (**Fig. 7C**), suggesting that increasing recurrent dominance is accompanied by reduced excitatory-to-inhibitory population activity. Next, to determine whether these changes were associated with altered system dynamics, we examined local dynamical stability and perturbation responses across diagnostic groups.

**Fig. 7.**
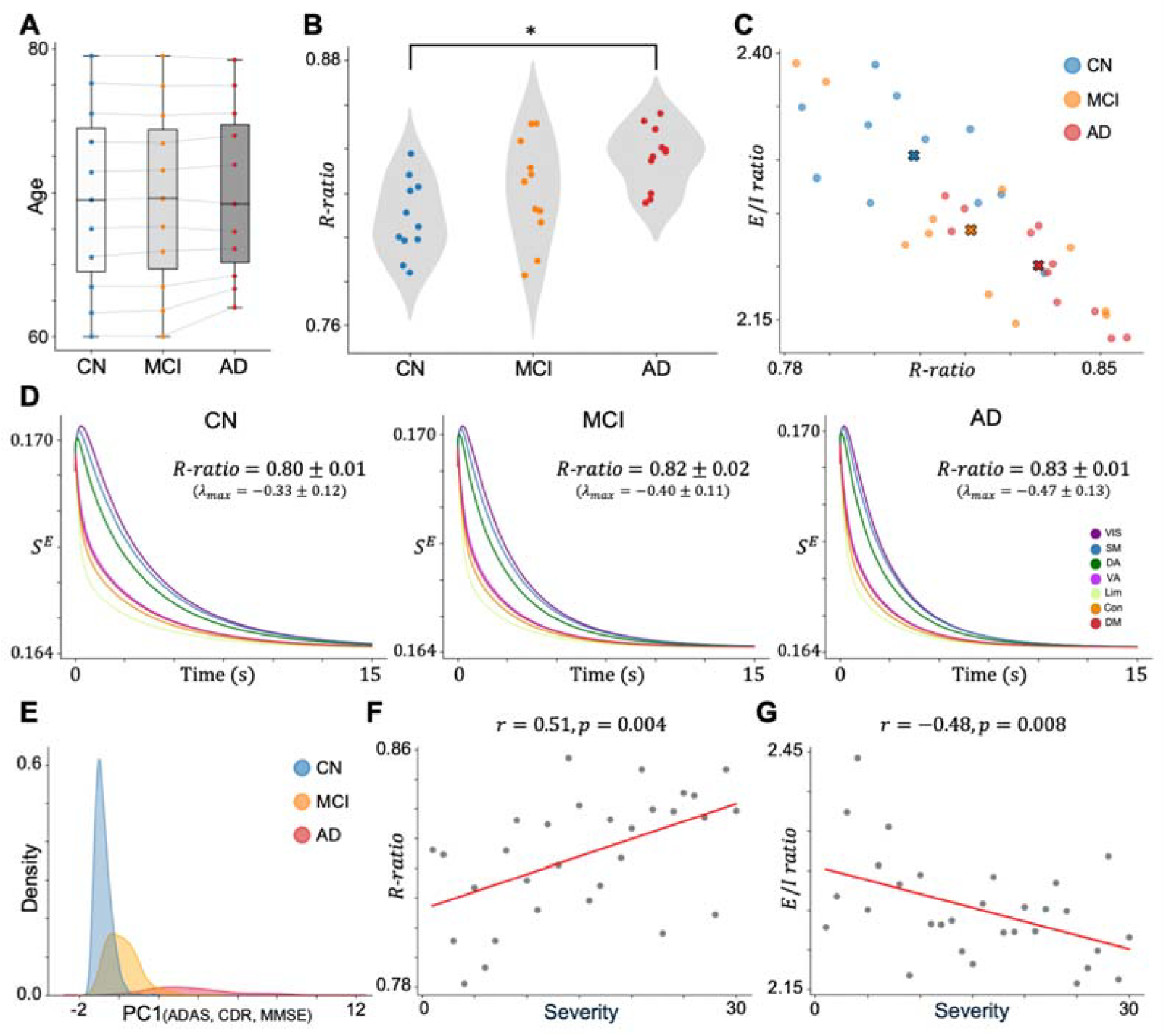
R-ratio alterations across the Alzheimer’s disease continuum. **(A)** Age distributions of the age-matched diagnostic groups. **(B)** Group differences in the R-ratio across CN, MCI, and AD groups. An asterisk indicates a statistically significant group difference (p < 0.05). **(C)** Distribution of the diagnostic groups according to the R-ratio and E/I ratio. Crosses indicate the group centroid. **(D)** Perturbation responses across the three diagnostic groups. Curves represent the mean excitatory synaptic gating activity averaged across fitted models within each group, with colors indicating intrinsic functional networks. **(E)** Distribution of participants along the continuous disease severity axis derived from PCA of clinical measures. Colors indicate diagnostic groups. **(F)** Relationship between disease severity and the R-ratio. **(G)** Relationship between disease severity and the E/I ratio. *Abbreviations: AD, Alzheimer’s disease; ADAS, Alzheimer’s disease assessment scale; CDR, clinical dementia rating; CN, cognitively normal; MCI, mild cognitive impairment; MMSE, mini-mental state examination; PCA, principal component analysis*.

Progressive decreases in local stability (λ_*max*_) and shorter activity persistence times following perturbation were observed from CN to AD (CN: 5.10 s ± 1.77 s; MCI: 3.75 s ± 0.80 s; AD: 3.28 s ± 0.80 s; **Fig. 7D**). These findings indicate that disease progression shifts the system farther from the bifurcation boundary and reduces the persistence of perturbation-induced activity. Finally, to characterize disease progression as a continuous process rather than discrete diagnostic categories, we examined trajectories of the R-ratio and E/I ratio along a continuous disease severity axis. Disease severity was quantified using the first principal component (PC1) from principal component analysis (PCA) of the Alzheimer’s disease assessment scale (ADAS), clinical dementia rating (CDR), and mini-mental state examination (MMSE) measurements, with higher PC1 values indicating greater disease severity (**Fig. 7E**). Along this continuous disease severity axis, the global R-ratio increased (r = 0.51, p = 0.004; **Fig. 7F**), whereas the global E/I ratio decreased progressively (r = −0.48, p = 0.008; **Fig. 7G**). Together, these findings demonstrate that disease progression across the AD continuum is characterized by a systematic increase in recurrent dominance, accompanied by reduced excitatory relative to inhibitory activity and diminished dynamical responsiveness.

## DISCUSSION

The balance between local recurrent and distributed inter-regional excitatory contributions is a fundamental principle underlying large-scale brain function, yet a quantitative framework for measuring this balance has been lacking. In the present study, we extended a biologically interpretable whole-brain circuit model to develop the R-ratio, a metric quantifying the relative balance between intra-regional recurrent excitation and inter-regional excitatory amplification. By jointly examining the R-ratio and the E/I ratio, we showed that these measures capture complementary aspects of cortical organization. Whereas the E/I ratio reflects the relative dominance of excitatory over inhibitory population activity, the R-ratio characterizes the source balance of excitatory drive. Using dynamical analyses, we further demonstrated that the R-ratio shapes whole-brain dynamical properties and responsiveness to perturbation. Finally, application of this framework to aging and AD revealed progressive increases in the R-ratio, suggesting that age- and disease-related brain alterations involve shifts in the balance between local recurrent and distributed inter-regional excitation. Together, these findings establish the R-ratio as a biologically interpretable framework for examining how the balance between intra- and inter-regional excitation shapes cortical organization, whole-brain dynamics, and brain health in aging.

Interactions within and between brain regions have long been recognized as fundamental components of large-scale brain dynamics, supporting local computation and distributed information integration. Previous studies have characterized these processes using FC-based network measures (Bassett and Gazzaniga, 2011; Bassett and Sporns, 2017; Sepulcre et al., 2010; Sporns, 2011), E/I ratio (Abeysuriya et al., 2018; Deco et al., 2014; Zhang et al., 2024), and functional gradients describing cortical organization (Margulies et al., 2016; Namgung et al., 2024; Taylor et al., 2026). Although these approaches provide valuable information regarding the strength and spatial organization of brain activity, they do not distinguish whether neural activity is predominantly driven by local recurrent excitation or distributed inter-regional excitatory contributions. The R-ratio complements these existing measures by quantifying the relative source of excitatory drive rather than its overall magnitude. Importantly, the R-ratio should not be interpreted as another measure of excitation or inhibition. Instead, it captures an complementary aspect of cortical dynamics: whether excitatory activity primarily arises from local or from distributed inter-regional interactions. Consistent with this interpretation, the R-ratio was systematically organized across intrinsic functional networks and aligned with MEG-derived temporal and spectral features. Higher R-ratio values were preferentially associated with regions exhibiting longer INT and slower oscillatory activity, consistent with computational studies showing that stronger recurrent excitation supports persistent activity, attractor dynamics, and information maintenance (Compte, 2000; Wang, 2002; Wong and Wang, 2006).

The biological interpretability of the R-ratio arises from both its model formulation and its correspondence with biological properties. The R-ratio was derived from a biologically interpretable large-scale circuit model that explicitly separates local recurrent excitation from long-range excitatory inputs, allowing direct quantification of their relative contributions. Cell-type-specific analyses revealed that astrocytes, microglia, oligodendrocyte precursors, pericytes, and endothelial cells were preferentially associated with cortical regions exhibiting low R-ratio values. These cell types contribute to synaptic homeostasis, immune regulation, myelination-related plasticity, and neurovascular coupling (De Faria et al., 2019; Guedes et al., 2022; Mahmoud et al., 2019; McConnell and Mishra, 2022), raising the possibility that low R-ratio regions are more strongly influenced by non-neuronal support systems. In contrast, excitatory neurons, inhibitory neurons, and oligodendrocytes were distributed along a linear axis within the joint R-ratio and E/I ratio space, suggesting coordinated relationships between recurrent processing, inhibitory regulation, and myelinated communication (Fields, 2015; Sadeh and Clopath, 2021; Seidl, 2014; Tatti et al., 2017). Together, these findings indicate that the R-ratio captures biologically meaningful organizational principles spanning cellular, structural, and functional levels.

Our dynamical analyses further demonstrated that the R-ratio governs whole-brain dynamical regimes. The optimal R-ratio was consistently located close to the bifurcation boundary, supporting the hypothesis that the human brain operates near a critical dynamical regime (Cocchi et al., 2017; Deco et al., 2013a; Luppi et al., 2026). Operating near this transition enables the system to maintain stable activity while remaining sufficiently sensitive to switch between dynamical states (Breakspear, 2017; Deco et al., 2017; Hancock et al., 2024). Perturbation analyses further showed that the optimal R-ratio exhibited prolonged persistence of stimulus-evoked activity while preserving stable return to the fixed point, indicating that stimulus-evoked activity could be maintained without diverging into unstable dynamics. Such intermediate dynamical regimes may facilitate both sustained information maintenance and efficient incorporation of new inputs (Breakspear, 2017; Rocha et al., 2022; Sanz Perl et al., 2022). Together, these findings suggest that the R-ratio provides a quantitative description of the balance between stability, flexibility, and responsiveness in whole-brain dynamics.

The R-ratio also provided new insights into age- and disease-related alterations in brain dynamics. Across both normal aging and the AD continuum, the R-ratio progressively increased, indicating a shift toward greater reliance on local recurrent excitation relative to distributed inter-regional input. This pattern is consistent with previous observations that long-range connections are particularly vulnerable to aging and AD, leading to reduced large-scale network integration (Liu et al., 2014; Sanz-Arigita et al., 2010; Tomasi and Volkow, 2012). Our findings therefore raise the possibility that age- and disease-related increases in the R-ratio reflect reduced contributions from distributed inter-regional communication rather than simple changes in overall neuronal activity. Moreover, progressive reductions in dynamical responsiveness across the AD continuum suggest that these alterations extend beyond static network organization and influence the dynamical operating regime of the brain. In this context, the R-ratio may provide a model-derived marker for quantifying age- and disease-related alterations in large-scale brain dynamics.

Several limitations should be considered. First, the R-ratio values may not be directly comparable across independent datasets because model parameters can be influenced by differences in imaging acquisition and preprocessing strategies. Future studies using harmonized multicenter datasets will be necessary for establishing the reproducibility and generalizability of the R-ratio across cohorts. Second, the present study focused on resting-state dynamics. Extending this framework to task-based or naturalistic functional MRI contexts may reveal how the balance between local recurrent and distributed inter-regional excitation is dynamically reconfigured under different cognitive demands (Cabalo et al., 2025; Finn and Bandettini, 2021; Hasson et al., 2004; Vanderwal et al., 2019; Wang et al., 2025). Third, although we focused on AD as a representative neurodegenerative disorder, further studies are needed to determine whether the R-ratio provides similar insights into other neurodegenerative and psychiatric disorders characterized by altered large-scale brain dynamics.

Overall, the present study establishes the R-ratio as a biologically interpretable computational framework for quantifying the balance between local recurrent and distributed inter-regional excitation. By linking local circuit-level excitatory balance mechanisms to large-scale brain dynamics, the R-ratio enables estimation of local circuit properties from macroscopic neuroimaging data. This framework may therefore allow future studies to track how the balance changes across development, aging, neurodegeneration, and psychiatric disorders. Moreover, it may provide a mechanistic basis for interpreting circuit-level reconfiguration associated with disease progression, cognitive state changes, and therapeutic interventions.

## METHODS

### Imaging data

#### i) HCP dataset

We analyzed multimodal MRI data of 1,004 neurologically healthy participants in the HCP-Young Adult dataset (Van Essen et al., 2013), which was used in the previous study (Zhang et al., 2024). T1-weighted MRI was acquired using a magnetization-prepared rapid gradient-echo (MPRAGE) sequence (repetition time (TR) = 2,400 ms, echo time (TE) = 2.14 ms, voxel resolution = 0.7 mm isotropic). T2-weighted MRI was acquired using a Sampling Perfection with Application-optimized Contrasts by using different flip angle Evolutions (SPACE) sequence (TR = 3,200 ms, TE = 565 ms, voxel size = 0.7 mm isotropic). Diffusion-weighted imaging (DWI) was acquired using a spin-echo echo-planar imaging (EPI) sequence (TR = 5,520 ms, TE = 89.5 ms, voxel resolution = 1.25 mm isotropic, diffusion weighting at b = 1000, 2000, and 3000 s/mm^2^, diffusion directions = 270, number of b0 images = 18). Resting-state functional MRI (rs-fMRI) was acquired using a gradient-echo EPI sequence (TR = 720 ms, TE = 33.1 ms, voxel resolution = 2 mm isotropic, number of volumes = 1,200).

#### ii) ADNI dataset

We analyzed 1,262 imaging sessions from the ADNI-GO, ADNI-2, and ADNI-3 datasets, including CN (N = 689), MCI (N = 436), and AD (N = 137) (Weiner et al., 2012). Only imaging sessions containing T1-weighted MRI, DWI, and rs-fMRI acquired from the same participant were included. For ADNI-GO and ADNI-2, T1-weighted MRI was acquired using an Inversion Recovery Spoiled Gradient Echo (IR-SPGR) sequence (TR = 6.98–7.65 ms, TE = minimum full echo time, voxel resolution = 1 mm isotropic). DWI was acquired using a spin-echo EPI sequence (TR = 9,050 ms, TE = minimum full echo time, diffusion weighting at b = 1000 s/mm^2^, diffusion directions = 41, number of b0 images = 5). Rs-fMRI was acquired using a gradient-echo EPI sequence (TR = 3,000 ms, TE = 30 ms, voxel resolution = 3.3 mm isotropic, number of volumes = 200). For ADNI-3, T1-weighted MRI was acquired using the MPRAGE sequence (TR = 2,300 ms, TE = minimum full echo time, voxel resolution = 1 mm isotropic). DWI was acquired using an EPI sequence (TR = 7,200 ms, TE = 56 ms, diffusion weighting at b = 1000 s/mm^2^, diffusion directions = 48, number of b0 images = 7 or TR = 3,300 ms, TE = 71 ms, diffusion weighting at b = 500, 1000, and 2000 s/mm^2^, diffusion directions = 112, number of b0 images = 15). Rs-fMRI was acquired using an EPI sequence (TR = 3,000 ms, TE = 30 ms, voxel resolution = 3.4 mm isotropic, number of volumes = 200). For rs-fMRI, only scans acquired with a TR of 3s and containing at least 193 volumes were retained. Because FCD analysis requires equal-length time series across participants, scans with more than 193 volumes were truncated to the last 193 time points.

### Preprocessing

#### i) HCP dataset

We used the same HCP dataset as in the previous study (https://github.com/ThomasYeoLab/CBIG/tree/master/stable_projects/fMRI_dynamics/Zhang_2024_pFIC/replication/HCP/input) (Zhang et al., 2024). Imaging data were preprocessed using the HCP minimal preprocessing pipeline (Glasser et al., 2013). T1w MRI data were corrected for gradient nonlinearity and bias field inhomogeneity. Non-brain tissues were removed, and the images were nonlinearly registered to the MNI152 standard space. Cortical white and pial surfaces were reconstructed from tissue boundaries, and mid-thickness and inflated surfaces were generated following established surface reconstruction procedures (Dale et al., 1999; Fischl, 2012; Fischl et al., 1999). The reconstructed surfaces were aligned to the fsLR surface space using multimodal surface matching and resampled to the 32k vertex mesh (Glasser et al., 2016; Robinson et al., 2014; Van Essen et al., 2012). DWI data were corrected for susceptibility-induced EPI distortions using reversed phase-encoding acquisitions, followed by eddy-current and head motion correction. Whole-brain streamline tractography was performed using the second-order integration over fiber orientation distributions (iFOD2) algorithm (Smith et al., 2012). Rs-fMRI data were corrected for EPI distortions and head motion, registered to the individual T1-weighted image, and subsequently transformed to MNI152 standard space. Structured noise components associated with head motion, white matter, cardiac pulsation, arterial signals, and large-vessel contributions were removed using ICA-FIX denoising (Salimi-Khorshidi et al., 2014). The preprocessed time series were high-pass filtered and mapped onto the standard fsLR grayordinate surface using cortical ribbon-constrained volume-to-surface mapping. The SC, FC, and FCD matrices were constructed using the Desikan–Killiany and Schaefer 100 atlases (Desikan et al., 2006; Schaefer et al., 2018). To construct the group-level SC matrix, a thresholding procedure was applied to reduce false-positive connections (Zhang et al., 2024). Specifically, connections with non-zero streamline counts in fewer than 50% of participants were removed from all individual SC matrices. For the remaining connections, streamline counts were averaged across participants with non-zero values and log-transformed to generate the group-level SC matrix. Individual FC matrices were computed by calculating Pearson correlations between regional rs-fMRI time series for each run, and then averaging the resulting matrices across runs. The group-level FC matrix was obtained by averaging individual FC matrices across participants. For FCD analysis, a sliding-window length of approximately 1 min (83 time points in the present dataset) was used, consistent with previous studies (Leonardi and Van De Ville, 2015; Liégeois et al., 2017). Using a step size of 1 TR, FC matrices were computed within each sliding window using Pearson correlations between regional time series. The upper triangular elements of each windowed FC matrix were vectorized, and pairwise correlations between these vectors were calculated to generate the FCD matrix. Group-level FCD distributions were obtained by averaging the cumulative distribution functions (CDFs) of individual FCD values across participants. For MP moments, T1w/T2w images were first aligned to the T1w image space. The cortical gray matter was divided into 14 equivolumetric surfaces, and T1w/T2w intensity values were sampled across cortical depth. Four moment features, including the mean, standard deviation, skewness, and kurtosis, were computed from each MP. Individual MP moments were then averaged across participants to obtain group-level MP moment maps.

#### ii) ADNI dataset

Imaging data were preprocessed using micapipe (Cruces et al., 2022). T1w MRI data were reoriented, corrected for intensity nonuniformity, skull-stripped, segmented into gray matter, white matter, and cerebrospinal fluid, and registered to the MNI152 template. Pial and white matter surfaces were reconstructed using FreeSurfer7 and subsequently registered to the fsLR surface space. DWI data were denoised using Marchenko-Pastur principal component analysis (MP-PCA), followed by Gibbs ringing correction and bias-field correction. Whole-brain tractography was performed using the iFOD2 algorithm, and streamline weights were estimated using spherical-deconvolution informed filtering of tractograms 2 (SIFT2) to reduce reconstruction biases. The atlas was then applied to generate individual SC matrices, and the group-level SC matrix was obtained by averaging individual SC matrices across participants. Rs-fMRI data were preprocessed after discarding the first five volumes. Head motion correction was then performed, followed by nuisance variable regression to remove confounding effects from white matter, cerebrospinal fluid, and head motion. FC and FCD were then computed from the preprocessed rs-fMRI time series using the same procedures as those applied to the HCP dataset. The only difference was that a sliding-window length of 20 TRs was used for FCD analysis, corresponding to an approximately 1-min window in the ADNI dataset.

### Conventional large-scale circuit modeling

Whole-brain blood-oxygen-level-dependent (BOLD) signals were simulated using the pFIC model, a neural mass model that extends the FIC framework by parameterizing local control variables to reduce the number of free parameters (Deco et al., 2013a; Zhang et al., 2024). Neural activity in the *i*^th^ cortical region was represented by coupled excitatory and inhibitory neuronal populations governed by the following nonlinear differential equations:

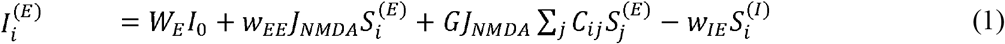

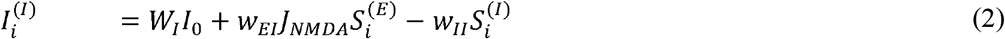

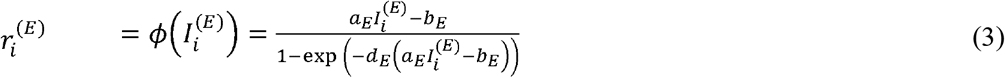

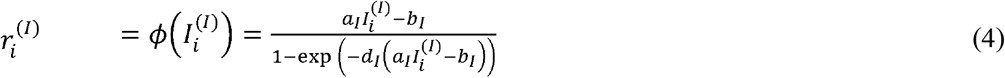

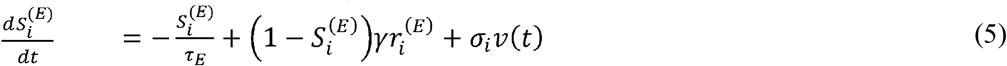

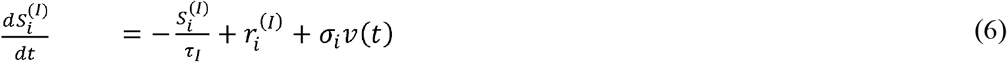

Here, *I, r*, and *S* denote synaptic input current, neuronal firing rate, and synaptic gating variable, respectively. The superscripts *E* and *I* denote excitatory and inhibitory neuronal populations, respectively. Equations (1) and (2) describe the synaptic input currents for the excitatory and inhibitory populations in the *i*^th^ cortical region. The term *W*_*x*_ *I* _o_, where x ∈{*E, I*} represents the external input current to population *x*. The parameter *w*_ab_ denotes the local connection strength from population *b* to *a*, where *a,b* {*E*, I}. J_NMDA_ represents the NMDA receptor-mediated synaptic coupling strength. Inter-regional excitatory input was mediated by the SC matrix *Cij*, which represents the connection strength from region *j* to region *i*, with its overall contribution scaled by a single global coupling parameter *G*. Following previous studies, *W*_*E*_, *W*_*I*_, *I*o, *w*_II_, and *J*_*NMDA*_ were set to 1, 0.7, 0.382 nA, 1, and 0.15 nA, respectively (Deco et al., 2013b; Zhang et al., 2024). Equations (3) and (4) transform synaptic input currents into population firing rates using nonlinear sigmoid transfer functions. Following previous studies, *a*_*E*_, *a*_*I*_, *b*_*E*_, *b*_*I*_, *d*_*E*_, and *d*_*I*_ were set to 310 n/C, 615 n/C, 125 Hz, 177 Hz, 0.16 s, and 0.087 s, respectively (Deco et al., 2013b; Zhang et al., 2024). Equations (5) and (6) describe the dynamics of the excitatory and inhibitory synaptic gating variables. The parameters *τ*_*E*_ and *τ*_*I*_ represent the excitatory and inhibitory synaptic time constants, respectively, and *γ* controls the saturation kinetics of synaptic activity. The term *v* denotes standard Gaussian noise scaled by the region-specific noise amplitude *σ*_*i*_.

The parameters *τ*_*E*_, *τ*_*I*_, andγ were set to 100 ms, 10 ms, and 0.641, respectively, following previous work (Deco et al., 2013b; Zhang et al., 2024). For each cortical region, the feedback inhibition parameter *w*_*IE*_ was computed such that the mean firing rate of the excitatory population remained close to 3 Hz during spontaneous activity (Demirtaş et al., 2019). This procedure maintains the model within a physiologically plausible operating regime while allowing regional differences in recurrent excitation, excitatory-to-inhibitory coupling, and stochastic fluctuations. Simulated excitatory synaptic gating variable was subsequently converted into BOLD signals using the Balloon–Windkessel hemodynamic model (Heinzle et al., 2016; Stephan et al., 2007).

The region-specific (*i*-th region) control parameters *wEE, wEI*, and σ_*i*_ are expressed as linear combinations of two cortical organizational features, including the principal gradient of resting-state FC and the T1w/T2w-derived cortical myelin estimate (Kong et al., 2021; Zhang et al., 2024):

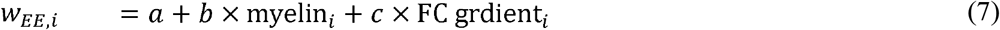

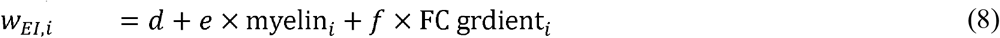

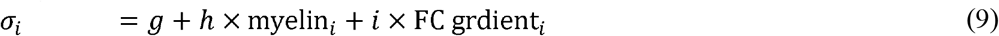

This parameterization reduces the number of trainable parameters to ten: nine coefficients for defining the three linear combinations and one scalar parameter, *G*. The ten trainable parameters were optimized by minimizing a loss function that quantified discrepancies between empirical and simulated static and dynamic FC. The static FC loss included the Pearson correlation coefficient (*r*) and L1 distance (*d*) between the vectorized upper triangles of the empirical and simulated FC matrices. The correlation term quantifies similarity in the overall spatial organization of FC, whereas the L1 term was included to prevent excessive synchronization in the simulated FC. The FCD loss was defined as the KS distance between the cumulative distributions of empirical and simulated FCD values. This term quantified discrepancies in the distribution of dynamic FC states between empirical and simulated FCD. The total loss was defined as (1 −*r*) + *d* + *KS*. Model parameters were optimized using the covariance matrix adaptation evolution strategy (Hansen, 2006). The HCP sample was divided into training (N = 335), validation (N = 335), and test (N = 334) datasets (Van Essen et al., 2013). Model parameters were estimated using the training dataset, model selection was performed using the validation dataset, and final performance was evaluated exclusively in the held-out test dataset.

### Modified biophysical model and derivation of recurrent-ratio

In the original pFIC model, recurrent excitation is parameterized through the local excitatory connection strength *w*_*EE*_, whereas inter-regional excitatory input is mediated by the SC uniformly scaled by a single global coupling parameter. To quantify the relative contribution of intra- and inter-regional excitatory inputs at the regional level, we modified the original pFIC formulation as follows:

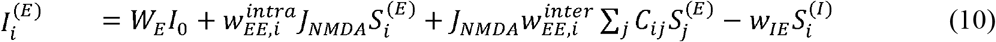

In the modified model, the original recurrent excitatory parameter *w*_*EE*_ was redefined as the intra-regional excitatory parameter 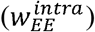. We further introduced a region-specific inter-regional excitatory parameter 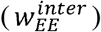 to quantify the gain of excitatory inputs transmitted through the SC. This formulation enabled intra- and inter-regional excitatory contributions to be estimated independently for each cortical region. The newly introduced parameter was modeled using the same linear combination strategy based on cortical myelin content and principal FC gradient (Glasser and Van Essen, 2011; Margulies et al., 2016):

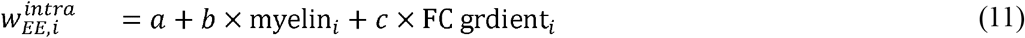

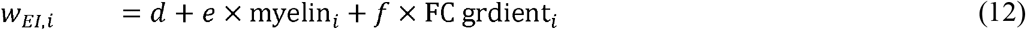

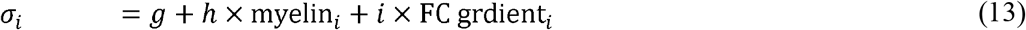

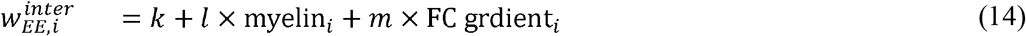

Because the ADNI dataset included substantially fewer participants than the HCP dataset (15 vs. 335 training sets), we incorporated spatial constraints during model fitting to improve parameter estimation. Following previous work that constrained *w*_*EI*_ to be negatively associated with myelin content and positively associated with the FC gradient (Zhang et al., 2024), we constrained 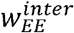 to exhibit the opposite spatial pattern, showing a positive association with myelin content and a negative association with the FC gradient. To characterize the spatial organization of the fitted parameters, we summarized 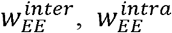, and *w*_*EI*_ across seven intrinsic functional networks, including visual, somatomotor, dorsal attention, ventral attention, limbic, frontoparietal, and default mode networks (Yeo et al., 2011). We further examined the relationship between 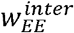 and cortical microstructure by calculating Pearson correlations between the optimal 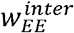 values and the HCP group-level MP moment features. Statistical significance was assessed using a spin permutation test implemented in the ENIGMA Toolbox (Larivière et al., 2021).

Based on the estimated intra- and inter-regional excitatory parameters, we defined the R-ratio of *i*-th region as follows:

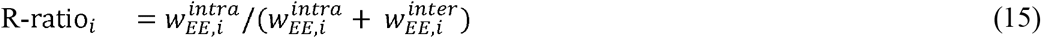

A higher R-ratio indicates that recurrent intra-regional excitation predominates over inter-regional excitatory input, suggesting that regional activity is primarily shaped by locally recurrent processing. Conversely, a lower R-ratio denotes stronger inter-regional excitatory influence, suggesting that regional activity is predominantly driven by distributed inputs from structurally connected brain regions. Thus, the R-ratio provides a quantitative measure of the relative dominance of local recurrent versus distributed excitatory processing.

### R-ratio and E/I ratio space analysis

The two-dimensional space defined by the R-ratio and E/I-ratio was constructed using the optimal control parameters. Following previous work, the optimal E/I ratio was estimated from simulations performed with these parameters (Zhang et al., 2024). Specifically, the regional E/I ratio was computed as the temporal average of the ratio between the excitatory and inhibitory synaptic gating variables,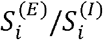. Each cortical region was represented as a point in the two-dimensional R-ratio and E/I ratio space and labeled according to its assignment to one of the seven intrinsic functional networks (Yeo et al., 2011). To assess how multimodal biological features were distributed within this space, we highlighted the top 10 cortical regions with the highest values for each multimodal feature. Functional features included HCP-derived MEG INT maps and band-specific power maps obtained through neuromaps (Markello et al., 2022; Van Essen et al., 2013). Structural features included HCP-derived cortical thickness and T1w/T2w ratio maps, also obtained through neuromaps (Markello et al., 2022; Van Essen et al., 2013). Cell-type-specific gene expression profiles were generated using regional gene expression data from the Allen Human Brain Atlas

(AHBA) processed with the abagen toolbox (Markello et al., 2021). Cell-type-specific gene sets were obtained from previous studies (Lake et al., 2018, 2016) and used to construct regional expression profiles for major brain cell classes. Eight representative cell types were examined, including astrocytes, endothelial cells, microglia, excitatory neurons, inhibitory neurons, OPCs, oligodendrocytes, and pericytes. For each cell type, regional expression intensity was computed by averaging the expression levels of the corresponding cell-type-specific genes within each cortical region.

### Dynamical responsiveness of the R-ratio

#### i) Parameter space construction

To examine how intra- and inter-regional excitatory inputs shape model dynamics, we constructed a two-dimensional parameter space by independently scaling the spatially varying components of 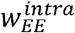 and 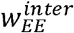:

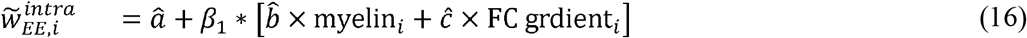

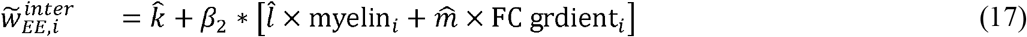

The hat notation indicates coefficients estimated from the optimally fitted model. The scaling factors *β*_1_and *β*_2_ were independently varied from 0 to 2 in increments of 0.1. This procedure modulated the regional heterogeneity of intra- and inter-regional excitatory inputs while preserving their optimized spatial patterns. For each pair of scaling factors, the scaled R-ratio was calculated as follows:

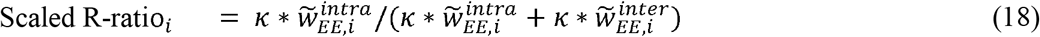

Because scaling 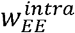 and 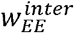 can shift the excitatory population away from its target firing rate of approximately 3 Hz, we introduced an additional normalization factor, *κ*_*i*_, to preserve the operating point of the excitatory population. The normalization factor *κ* was calculated as follows:

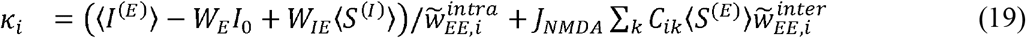

Here, ⟨⋅⟩ denotes the steady-state value of the corresponding variable required to maintain the excitatory firing rate, 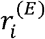, at approximately 3 Hz. The normalization rescales the overall magnitude of excitatory input while preserving the relative balance between intra- and inter-regional excitatory processing represented by the scaled R-ratio.

#### ii) Identification of bifurcation boundary

To determine whether changes in the relative magnitude of intra- and inter-regional excitatory inputs alter the dynamical regime of the model between stable and unstable systems, we performed a bifurcation analysis. For each pair of scaling factors (*β*_1_, *β*_2_), we calculated the fixed point of the deterministic system, which represents the equilibrium state around which spontaneous neural activity evolves. To assess the local stability near the fixed point, we linearized the deterministic system around the fixed point and constructed the corresponding Jacobian matrix, whose elements are given by the partial derivatives of the dynamical equations as follows:

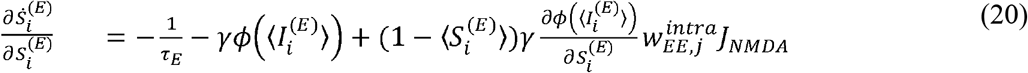

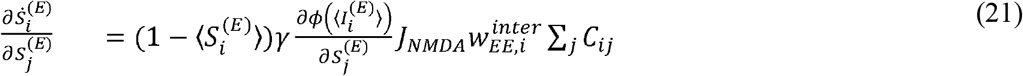

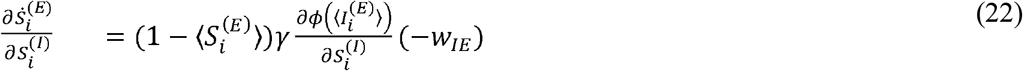

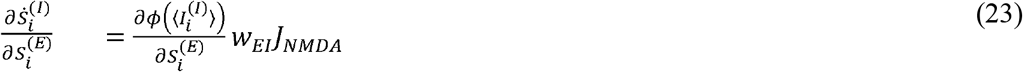

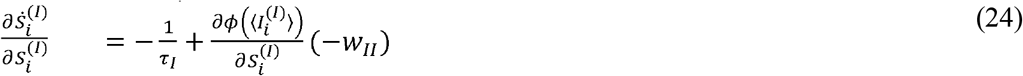

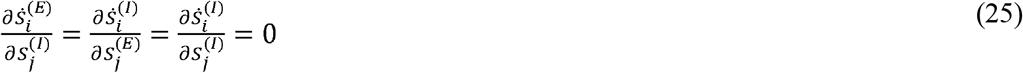

Here, ⟨⋅⟩ denotes the steady-state value of each variable evaluated at the fixed point. The Jacobian matrix characterizes the local response of the system to small perturbations around the fixed point and therefore determines whether the equilibrium is dynamically stable. To assess the stability of each fixed point, we computed the eigenvalues of the Jacobian matrix. Because the model contains both excitatory and inhibitory synaptic gating variables for every cortical region, the number of eigenvalues was twice the number of cortical regions. Local stability was determined by the largest real part of the eigenvalues, *λ*_max_ =max (Re(*λ*)). If *λ*_max_ < 0, all perturbations decay over time, and the fixed point is locally stable. Conversely, if λ_max_ > 0, perturbations grow exponentially, and the fixed point becomes unstable. The bifurcation boundary was therefore defined as the contour satisfying *λ*_max_ = 0, which separates stable and unstable dynamical regimes within the (*β*_1_, *β*_2_ parameter space. This analysis allowed us to determine how the relative balance between intra- and inter-regional excitation governs transitions in whole-brain dynamical stability.

#### iii) Intrinsic dynamics analysis

To characterize the intrinsic dynamical properties of varying R-ratio values, we quantified two complementary measures derived from simulations: (i) the E/I ratio and (ii) the STD-KOP. The E/I ratio was used to quantify the relative activity of excitatory and inhibitory populations. It was calculated as the ratio of excitatory and inhibitory synaptic gating variables, *S*^(*E*)^ and *S*^(*I*)^, obtained from spontaneous simulations. In contrast to the R-ratio, which quantifies the relative contributions of local recurrent versus inter-regional excitatory inputs, the E/I ratio reflects the relative dominance of excitatory over inhibitory population activity. The STD-KOP was used to quantify temporal fluctuations in global synchronization during spontaneous activity. The KOP measures the degree of phase synchronization across cortical regions at each time point, whereas its temporal standard deviation (i.e., STD-KOP) quantifies the variability of global synchronization over time. STD-KOP is widely used as a proxy for metastability, with higher values indicating more frequent transitions between synchronized and desynchronized states and lower values indicating more stable synchronization dynamics (Hancock et al., 2024; Kuramoto, 1984). To calculate STD-KOP, simulated BOLD signals were band-pass filtered between 0.01 and 0.08 Hz using a fourth-order Butterworth filter. The instantaneous phase of each regional BOLD signal was then extracted using the Hilbert transform. At each time point, the KOP was computed from the phases across all cortical regions, and STD-KOP was defined as the temporal standard deviation of the KOP over the entire simulation.

#### iv) Perturbation response analysis

To characterize the responsiveness and persistence dynamics associated with different R-ratio values across the parameter space, we performed a perturbation response analysis. For each combination of scaling parameters, the deterministic system was first initialized at its stable fixed point. A small perturbation (δ) was then applied to the synaptic gating variables to transiently displace the system from equilibrium. Specifically, perturbations were introduced by adding a small positive offset (+ δ o 0.005) to the excitatory synaptic gating variable or a small negative offset (−δ = −o 0.01) to the inhibitory synaptic gating variable.

Following the perturbation, the system was numerically integrated with a time step of dt (0.0005) to track its return toward the equilibrium state. Simulation durations were 27.5 s for the HCP analysis and 15 s for the ADNI analysis. For stable regimes in which the excitatory synaptic gating variable returned to the fixed point after perturbation, activity persistence time was defined as the time required for the mean excitatory activity to recover 80% toward the fixed point.

### Aging-related differences in the R-ratio

We examined age-related differences in the R-ratio using multiple imaging sessions of CN participants (N = 689) from the ADNI dataset (Weiner et al., 2012). Participants were sorted by age and grouped into consecutive age bins of 30 individuals. Within each age group, participants were randomly divided into training and test sets, with 15 participants assigned to each set. The parameter set yielding the lowest test loss was selected to estimate the group-level R-ratio. We then examined whether age-related differences in the R-ratio were associated with amyloid burden, cortical thickness, brain volume, and neurocognitive performance. Amyloid burden was quantified using amyloid positron emission tomography standardized uptake value ratio (SUVR) obtained from the Dallas Lifespan Brain Study (DLBS) dataset (Park et al., 2025). For each R-ratio age group, amyloid SUVRs were averaged across participants within an age window of ±2 years centered on the median age of the corresponding group. Cortical thickness was estimated from the CN participants in ADNI and averaged using the same age groups defined for the R-ratio analysis. Gray matter, white matter, and ventricular volumes were obtained from the Lifespan BrainChart Project (Bethlehem et al., 2022), and the median values corresponding to each age group were extracted. Neurocognitive performance was evaluated using the NeuroCognitive Performance Test (NCPT), a web-based cognitive assessment comprising multiple tasks (Jaffe et al., 2022). Individual tests were grouped into four cognitive domains: (i) executive function (go/no-go and Trail Making A/B tasks), (ii) memory (forward memory span, reverse memory span, and complex memory span), (iii) visual attention (divided visual attention, Posner cueing, dual search, and object recognition), and (iv) reasoning (arithmetic reasoning, grammatical reasoning, progressive matrices, scale balance, and digit symbol coding). For each R-ratio age group, cognitive scores were averaged across participants within an age window of ±2 years. Subtests were excluded if responses were available from fewer than 2,000 participants within the corresponding age window. All cognitive scores were adjusted such that higher values indicated poorer cognitive performance.

### Characteristics of the R-ratio across the Alzheimer’s disease continuum

We examined differences in the R-ratio across the AD continuum, including CN, MCI, and AD, using data from the ADNI dataset (Weiner et al., 2012). To minimize the influence of aging independent of disease progression, participants were age-matched across diagnostic groups. Specifically, age-matched groups were constructed in 2-year intervals between 60 and 80 years by selecting 30 participants from each diagnostic group within each age interval. Each group was divided into training and test sets, with 15 participants assigned to each set, and the parameter set yielding the lowest test loss was used to estimate the group-level R-ratio. Next, we examined how model dynamics change across disease progression. To this end, the mean R-ratio and *λ*_*max*_ were calculated for each age-matched group. Here, *λ*_*max*_ was defined as the maximum real part of the Jacobian eigenvalues evaluated at the fixed point, which quantifies the local dynamical stability of the system. To further investigate how disease-related alterations influence dynamical responsiveness, external perturbations were applied to the system. The activity persistence time was defined as the time required for the mean excitatory activity to recover 80% toward the fixed-point following perturbation. We further examined whether the R-ratio and E/I ratio varied continuously with disease severity. A composite disease severity score was derived from the ADAS, CDR, and MMSE. To ensure that higher values consistently reflect greater disease severity, MMSE scores were reverse-coded prior to analysis. PCA was then applied to these three scores, and PC1 was used as a continuous disease-severity axis. Participants were ranked according to their PC1 scores and grouped into consecutive sets of 30 individuals. Each group was subsequently divided into training and test sets, with 15 participants assigned to each set. For each severity group, the parameter set yielding the lowest test loss was selected to estimate the corresponding group-level R-ratio and E/I ratio.

## DATA AVAILABILITY

Imaging and phenotypic data were provided in part by the HCP (https://www.humanconnectome.org/; https://github.com/ThomasYeoLab/CBIG/tree/master/stable_projects/fMRI_dynamics/Zhang2024_pFIChang2024_pFIC), ADNI (https://adni.loni.usc.edu), neuromaps (https://github.com/netneurolab/neuromaps), abagen (https://github.com/rmarkello/abagen), DLBS (https://openneuro.org/datasets/ds004856/versions/1.3.0), Lifespan BrainChart Project (https://github.com/brainchart/Lifespan), and NCPT (https://github.com/pauljaffe/lumos-ncpt-tools/tree/v1.1.0).

## CODE AVAILABILITY

The code for the R-ratio analysis is available at https://github.com/CAMIN-neuro/Rratio.

## AUTHOR CONTRIBUTIONS

Y.P. and B.P. designed the study, analyzed the data, and wrote the manuscript. S.K. aided experiments. H.P, T.D.S., and B.C.B. reviewed the manuscript. B.P. is the corresponding author of this study and is responsible for the integrity of data analysis. All authors reviewed and approved the final version of the manuscript for publication.

## Supplementary Information

**Supplementary Fig. 1.**
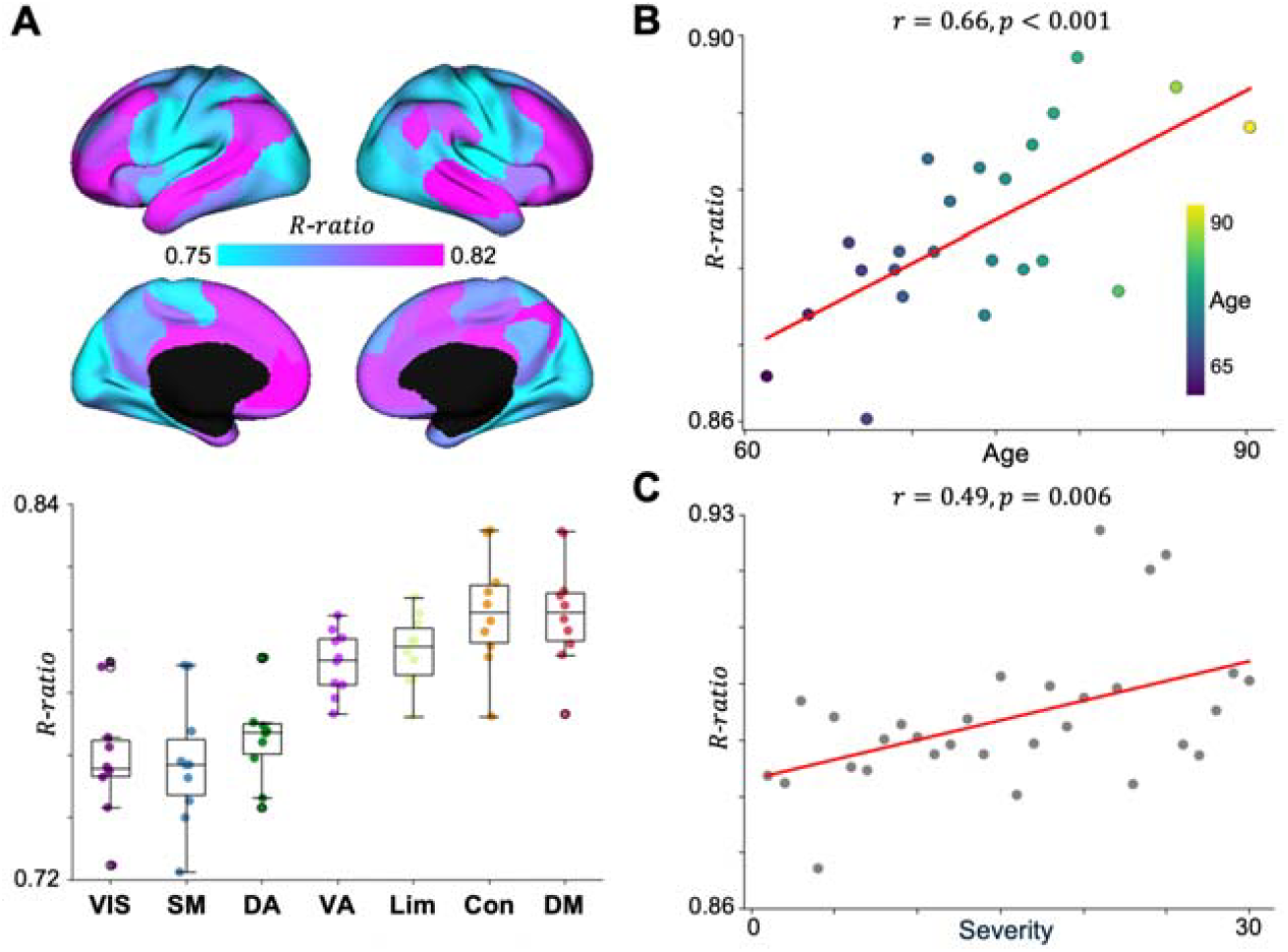
Control analysis using the Schaefer 100 atlas. **(A)** Spatial distribution of the R-ratio is shown on cortical surfaces, and the values are stratified across intrinsic functional networks (bottom). Box plots show the interquartile range, with center lines indicating median values. **(B)** Age-related differences in the R-ratio. **(C)** Relationship between disease severity and the R-ratio. *Abbreviations: VIS, visual; SM, somatomotor; DA, dorsal attention; VA, ventral attention; Lim, limbic; Con, frontoparietal control; DM, default mode; R-ratio, recurrent-ratio*.

**Supplementary Fig. 2.**
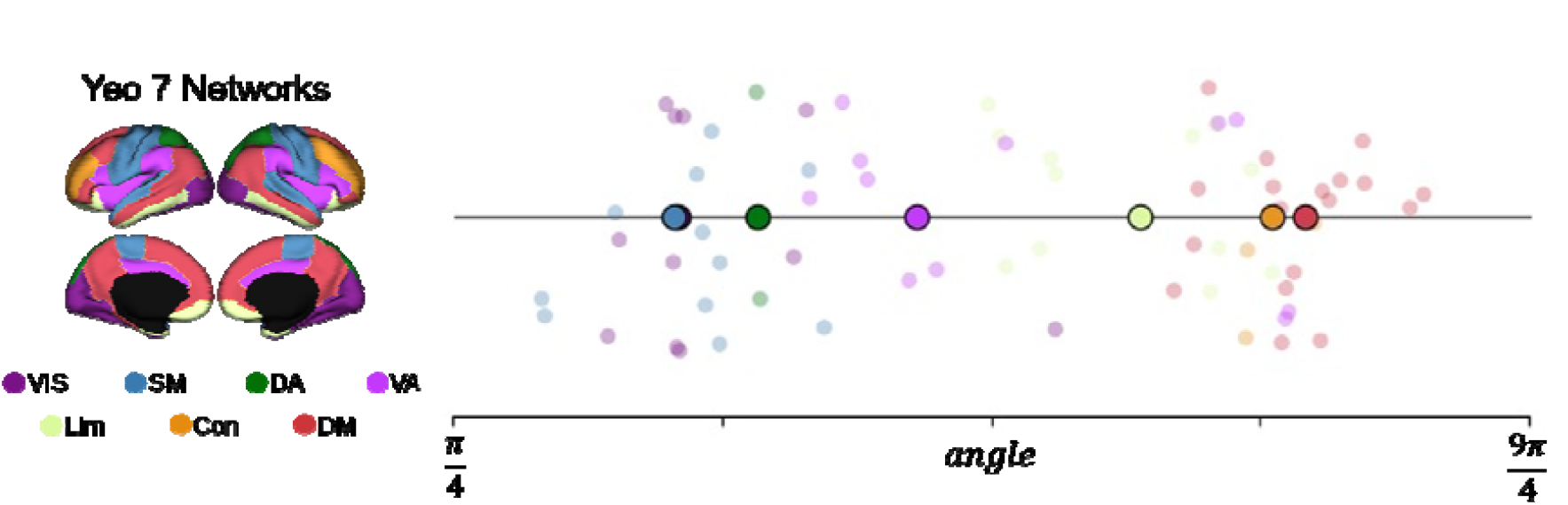
Functional networks are organized along the angular axis of the R-ratio and E/I ratio space. Angles were computed from z-scored R-ratio and E/I ratio values relative to the whole-cortex mean point. Faded markers indicate individual cortical regions, and enlarged markers indicate the centroid of each network along the angular axis. *Abbreviations: VIS, visual; SM, somatomotor; DA, dorsal attention; VA, ventral attention; Lim, limbic; Con, frontoparietal control; DM, default mode; R-ratio, recurrent-ratio*.

**Supplementary Fig. 3.**
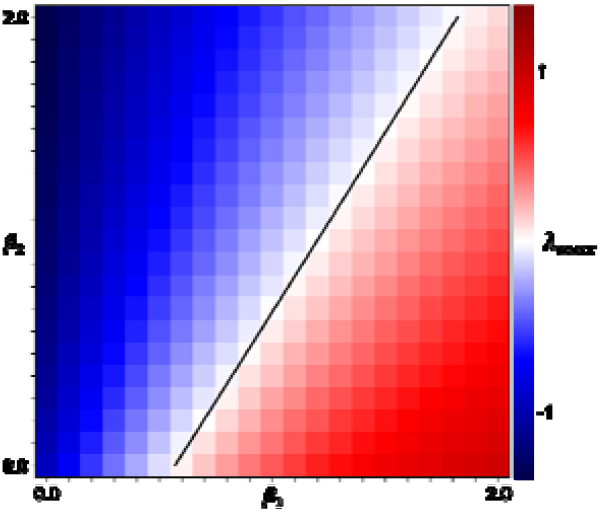
Stability landscape and bifurcation boundary of the scaled R-ratio within the parameter space. The color map shows the maximum real part of the Jacobian eigenvalues () evaluated at the fixed point across combinations of the scaled intra- and inter-regional excitatory variables. The black line indicates the bifurcation boundary (), which separates stable () and unstable () dynamical regimes.

**Supplementary Fig. 4.**
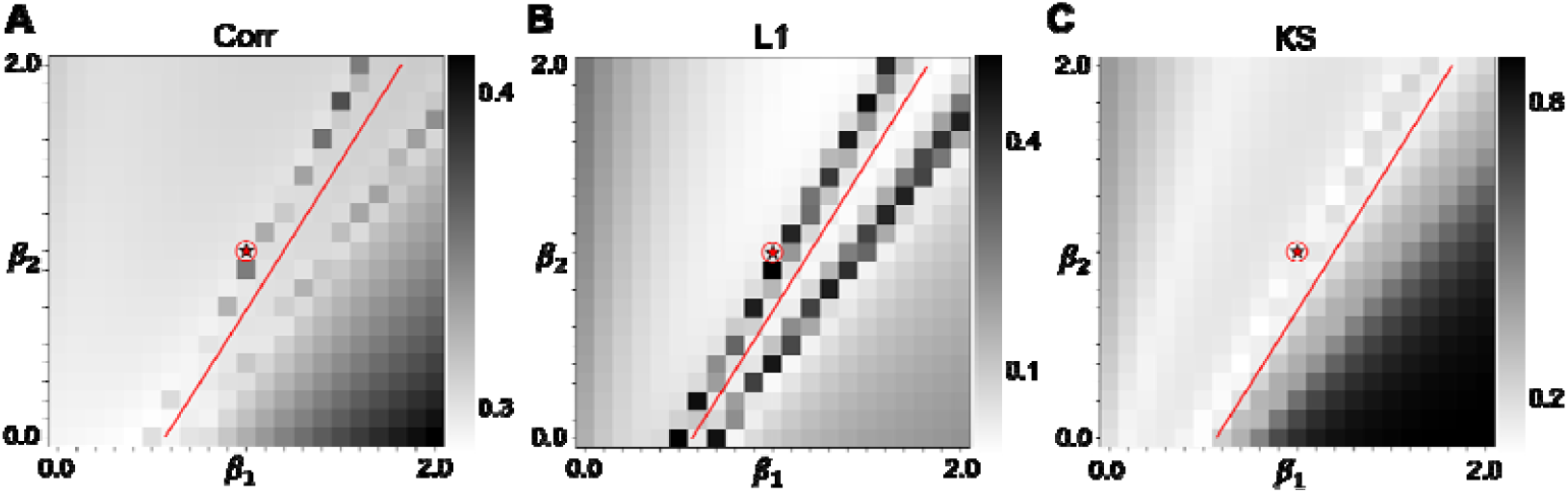
Individual loss components across the scaled R-ratio parameter space. **(A)** Pearson correlation, **(B)** L1 distance, and **(C)** KS statistics are shown across the parameter space. The red circle with a star indicates the optimal parameter combination, and the red line denotes the bifurcation boundary separating stable and unstable dynamical regimes. *Abbreviations: Corr, Pearson correlation; KS, Kolmogorov–Smirnov*.

